# Automated segmentation and quantitative analysis of organelle morphology, localization and content using CellProfiler

**DOI:** 10.1101/2022.11.09.515818

**Authors:** Sebastiaan N.J. Laan, Richard J. Dirven, Jeroen Eikenboom, Ruben Bierings, for the SYMPHONY consortium

**Author notes:** Corresponding author: Dr. Ruben Bierings, Department of Hematology, Erasmus University Medical Center, Rotterdam, The Netherlands.

## Abstract

One of the most used and versatile methods to study number, dimensions, content and localization of secretory organelles is confocal microscopy analysis. However, considerable heterogeneity exists in the number, size and shape of secretory organelles that can be present in the cell. One thus needs to analyze large numbers of organelles for valid quantification. Properly evaluating these parameters requires an automated, unbiased method to process and quantitatively analyze microscopy data. Here, we describe two pipelines, run by CellProfiler software, called OrganelleProfiler and OrganelleContentProfiler. These pipelines were used on confocal images of endothelial colony forming cells (ECFC) which contain unique secretory organelles called Weibel-Palade bodies. Results show that the pipelines can quantify the cell count and size, and the organelle count, size, shape, relation to cells and nuclei, and distance to these objects. Furthermore, the pipeline is able to quantify secondary signals located in or on the organelle or in the cytoplasm. Cell profiler measurements were checked for validity using Fiji. To conclude, these pipelines provide a powerful, high-processing quantitative tool for analysis of cell and organelle characteristics. These pipelines are freely available and easily editable for use on different cell types or organelles.

## Introduction

Eukaryotic cells are compartmentalized into organelles, subcellular entities separated from the cytoplasm by a limiting membrane that enable them to more efficiently carry out specialized functions in the cell, such as energy production and protein synthesis, transport and degradation. A specific class of organelles consists of secretory vesicles, which serve to temporarily store and then rapidly secrete molecules into the extracellular space on demand. Secretory organelles are vital to maintaining homeostasis, as they allow a cell to communicate with other, distant cells or to respond to immediate changes in its environment, such as in the case of injury or when encountering pathogens. Their function is often defined by the content that is secreted, which is cell type and context specific, and depends on a sufficient magnitude of release, which directly relates to the number and dimensions of the secretory organelles that can undergo exocytosis. Moreover, the intracellular location of secretory organelles in relation to their site of biogenesis (i.e. the Golgi apparatus), filaments of the cytoskeleton and the plasma membrane also indirectly determines their exocytotic behavior.

Weibel-Palade bodies (WPBs) are cigar-shaped endothelial cell specific secretory organelles that contain a cocktail of vasoactive molecules that are released into the circulation in response to vascular injury or stress (1). WPBs owe their typical elongated morphology to the condensation of its main cargo protein, the hemostatic protein Von Willebrand factor (VWF), into parallel organized tubules that unfurl into long platelet-adhesive strings upon release (2). The size and shape of WPBs are of interest from a biological and medical perspective as they correlate with the hemostatic activity of the VWF strings that are released (3) and can be reflective of disease states, such as in the bleeding disorder Von Willebrand Disease (VWD) (4). A model frequently used to study the pathophysiology of vascular diseases like VWD is the Endothelial Colony Forming Cell (ECFC). A major advantage of this model is that ECFCs can be derived from whole blood of patients, which allows analysis of patient endothelial cell function, WPB morphology and secretion *ex vivo*. However, substantial phenotypic heterogeneity can exist between ECFCs (5, 6), which stresses the need for robust quantitative analytical methods to evaluate their phenotype.

One of the most used and versatile methods to study number, dimensions, content and localization of secretory organelles is confocal microscopy analysis. However, as with all biological samples, considerable variability exists in the number, size and shapes of secretory organelles that can be present in the cell. One thus needs to analyze large numbers of organelles while ideally collecting this information in such a manner that it can be analyzed in a cell-by-cell manner. The crowded intracellular environment in combination with optical and immunostaining limitations presents an additional, technical challenge to separate individual organelles, which often precludes analysis on single organelle detail. Proper evaluation of these parameters requires an automated, unbiased method to process and quantitatively analyze microscopy data.

Here we describe 2 pipelines developed in CellProfiler (7), a free, easy to use image analysis software that uses separate module-based programming, for the identification, quantification and morphological analysis of secretory organelles within endothelial cells. The automated analysis pipeline OrganelleProfiler (OP) segments cells, WPBs, nuclei and cell membranes from microscopy images, quantifies number, location, size and shape of WPBs and extracts these data per cell and relative to the location of nucleus and perimeter of the cell. The function of the OrganelleProfiler pipeline is demonstrated by automated analysis of 2 previously established phenotypic classes of healthy donor ECFCs (6), which identifies clear differences in number, length, eccentricity and intracellular localization of WPBs. A second pipeline, called OrganelleContentProfiler (OCP), expands on the capabilities of the OrganelleProfiler by offering additional modules to measure the intensity of proteins of the secretory pathway both inside and outside the WPB. As an example we analyzed the presence of the small GTPase Rab27A on WPBs (8–10) and used protein disulfide isomerase (PDI), a marker for the endoplasmic reticulum, as a control as this marker is not present on WPBs.

Our CellProfiler pipelines provide robust and unbiased quantitative analysis tools for WPB morphometrics and can, with minimal adaptation, also be used to obtain quantitative data for other organelles and/or other cellular systems.

## Materials & Methods

### Endothelial Colony Forming Cells and Ethical Approval

General cell culture of Endothelial Colony Forming Cells (ECFCs) was performed as described (5). The study protocol for acquisition of the ECFCs was approved by the Leiden University Medical Center ethics review board. Informed consent was obtained from three subjects in accordance with the Declaration of Helsinki. Healthy participants were 18 years or older and had not been diagnosed with or known to have VWD or any other bleeding disorder. The ECFCs used in this study have previously been classified as group 1 or group 3 (6).

### Immunofluorescence of ECFCs and Image Acquisition

In short, ECFCs were grown on glass coverslips (9mm) and left confluent for 5 days before fixing with 70% methanol on ice for 10 minutes. Samples for OrganelleProfiler were stained with antibodies against VWF and VE-cadherin and nuclei were stained with Hoechst (S1 table for supporting information on antibodies). Samples for OrganelleContentProfiler were stained with Hoechst and antibodies against VWF, VE-cadherin and either Rab27A or PDI. After staining with appropriate fluorescently labeled secondary antibodies, coverslips were mounted using ProLong^®^ Diamond Antifade Mountant (Thermo Fisher Scientific). Visualization of the cells for the OrganelleProfiler example was done using the Imagexpress Micro Confocal System using the 63x objective without magnification. The OrganelleContentProfiler samples were imaged using the Zeiss LSM900 Airyscan2 upright confocal microscope using the 63x oil immersion objective. For both the OrganelleProfiler and OrganelleContentProfiler images a Z-stack was made which was transformed to a maximum Z-projection.

### CellProfiler-Based pipelines for cell organelle analysis and manual scoring with Fiji

CellProfiler (version 4.2.1 at time of publication) was used, which can be downloaded from the CellProfiler website (11). Images have to be of high enough resolution that individual organelles can be identified and do not blur together. Magnification, laser intensity, detector sensitivity and other acquisition parameters should be the same for each image. Image format has to be similar as well. We recommend uncompromised TIFF files. Pipelines developed are available in the Supplementary Materials (S1 and S2 file). To compare the CellProfiler measurements and validate these, manual scoring of cell count, cell surface area, WPB count, WPB length, and VWF and Rab27A intensity inside and outside the WPBs was performed using Fiji version 2.3.0 (12). Scoring was performed by using the build in scale and drawing regions of interest per cell and per WPB.

### Statistical Analysis

Output data of the OrganelleProfiler pipeline was compared by Mann-Whitney test if not normally distributed data and unpaired T test with Welch’s correction was performed on normally distributed data. Data of the OrganelleContentProfiler pipeline was compared with RM one way ANOVA with Geisser-Greenhouse correction. Data are presented as median with min/max boxplot. Results with p value < 0.05 were considered statistically significant. P values are indicated on the graphs in the figures. Data analyses was performed using GraphPad Prism 9.3.1 (GraphPad Software, San Diego, CA, USA).

## Results

### OrganelleProfiler (OP) - Automated identification and quantification of nuclei, cells and secretory organelles

Described here are the modules used in the OrganelleProfiler pipeline for the identification and measurement of endothelial cells, their nuclei and WPBs. The most important parameters and how these can be adjusted for use on other tissues for each module are mentioned in S3 file. Full explanations of other variables are available from the help function within the CellProfiler software or from the user manual on the CellProfiler website (11). For the development of OrganelleProfiler we used confocal images from ECFC clones from several healthy donors. These ECFCs have previously been classified into separate phenotypic groups based on cellular morphology and showed clear differences in expression of cell surface markers, proliferation and storage and secretion of VWF (6). Representative images of group 1 (top) and group 3 (bottom) ECFCs used for this study are shown in Fig 1. The CellProfiler modules that together form the OrganelleProfiler pipeline can be divided into 6 steps (Fig 2), which are described below.

**Fig 1.**
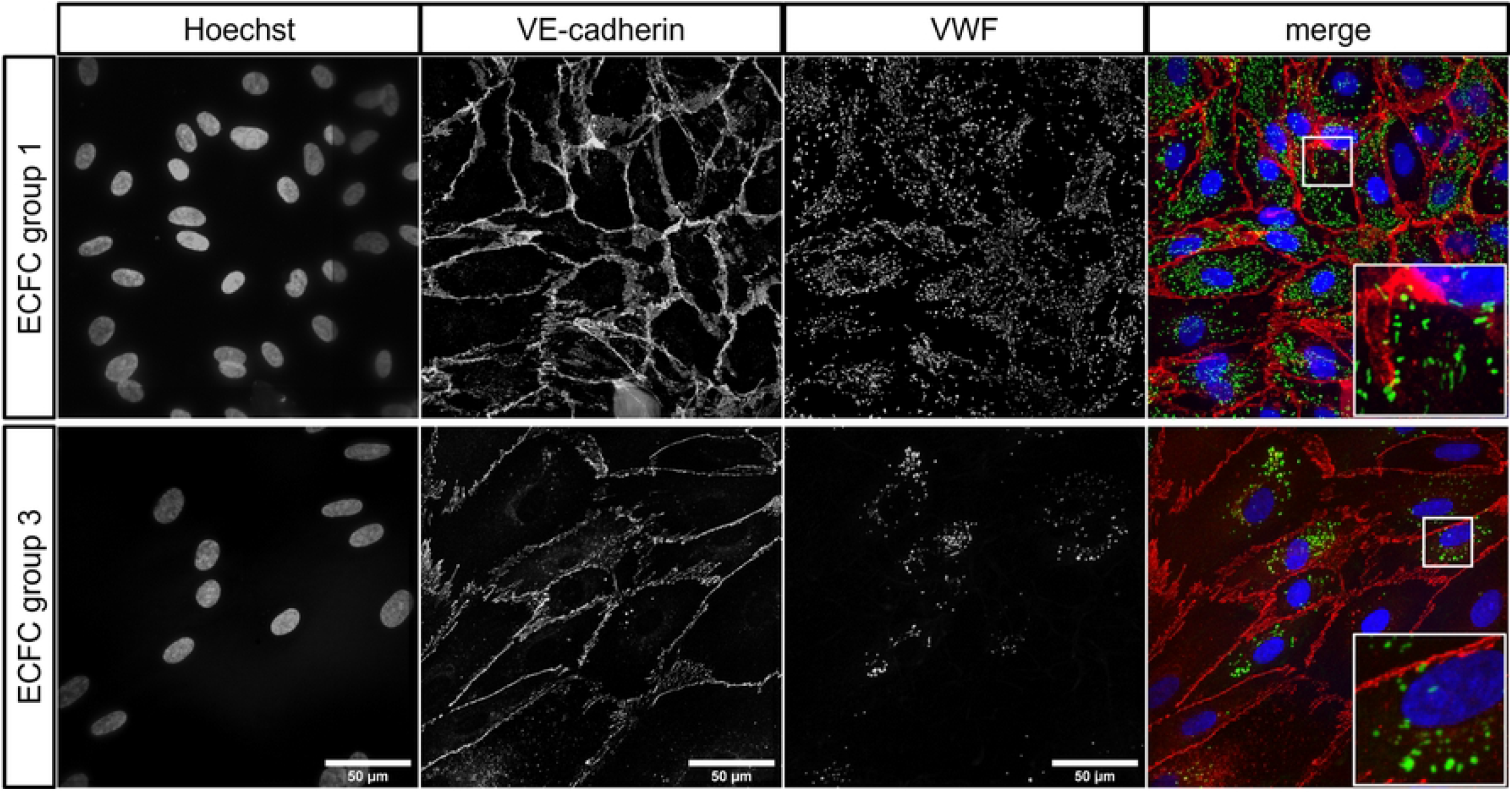
Representative images of healthy ECFC controls belonging to previously classified groups based on morphology (6). Group 1 ECFCs (top) and group 3 ECFCs (bottom) were stained with Hoechst (blue) and antibodies against VE-cadherin (red) and VWF (green). Scale bar represents 50 μm. Images were taken with a 63x objective.

**Fig 2.**
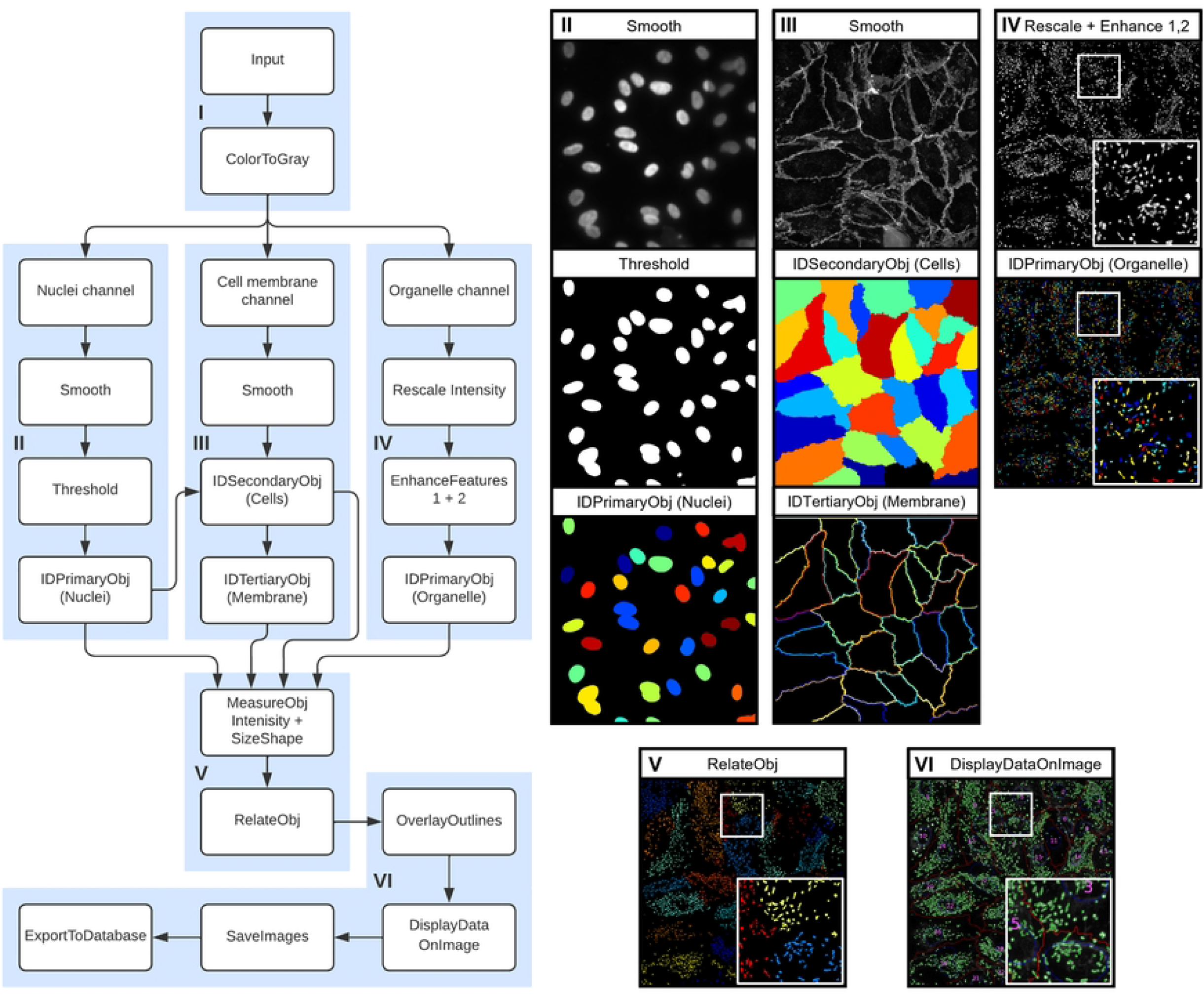
OrganelleProfiler: Quantitative and qualitative analysis of cells and cell secretory organelles. Left, flowchart of the modules within the OrganelleProfiler pipeline. I) Input of images and splitting of channels. II) Smoothing (top), thresholding (middle) and identification of the nuclei (bottom). Every different color indicates a different object. III) Smoothing of the cell membrane (top), identification of the cells (middle) and identification of cell membranes (bottom) as objects. IV) VWF signal rescaling and enhancement (top) and identification of WPB objects (bottom). V) Relating WPBs and Cells as child and parent respectively. Same colored objects indicate a relationship to the same cell. VI) Generated output image overlaying the outline of the nuclei (blue), cells (red), and WPBs (green) objects on the VWF channel. With the addition of the cell number (purple).

#### Step I – Input of images

Firstly, images of interest are imported into the software. In this example, 5 images from two groups of ECFCs were compared. Each image has 3 channels, 1 for the nuclei staining (Hoechst), one for cell membrane staining (VE-cadherin) and a third channel for organelle specific staining (VWF) (Fig 1). Channels are separated at this point so that each channel is processed separately in the following steps.

#### Step II, III and IV – Identification of nuclei, cell membranes, cells and organelles

Second, the nuclei staining signal is smoothed and a threshold is applied for the identification of the nuclei as objects. This object, together with the smoothed cell membrane staining signal is used in step III for the identification of the whole cell as secondary object. The nuclei are used as a starting point from which the object propagates outward in all directions until it encounters a secondary signal, in this case the smoothed cell membrane. A third object is generated using the cell object. This third object consists of only the cell membrane which is needed in the OrgannelleContentPipeline. In parallel to steps II and III, step IV uses the organelle staining signal for identification of the organelles. The signal is first rescaled and the speckle and neurite features are enhanced, which yields a better separation of organelles if they are located close to, or on top of, each other. After modification, the organelles are identified as the fourth object class.

#### Step V – Measurement and relating of objects

All objects that are generated in step II, III and IV are measured here. Size, shape and intensity, where relevant, is measured. Organelle objects are related to the nuclei and to the cell membrane in this step as well. This yields counts of secondary objects (organelles) per primary objects (cells) and distance of the secondary object to either the nuclei or the cell membrane. Measurements that we performed on the objects are eccentricity (as indicator for round or elongated WPB morphology), length of WPBs (maximum ferret diameter) and absolute as well as relative distance of WPBs to the nuclei and the cell membrane (Fig 3A).

**Fig 3.**
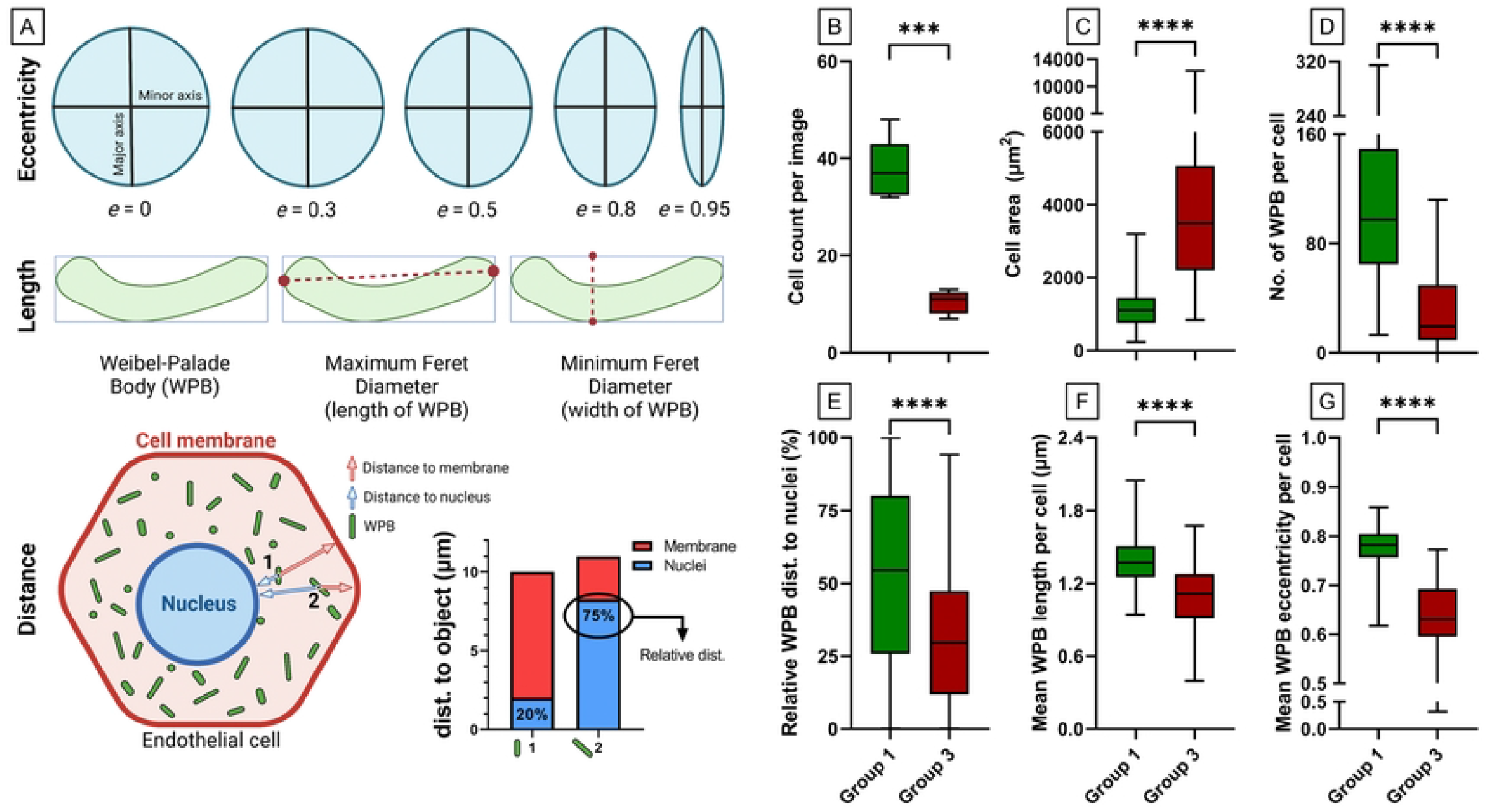
Quantitative and morphological differences between ECFC control groups. Two **previously classified ECFCs based on morphology (6)**, group 1 (green) and group 3 (red), were stained for Hoechst, VE-cadherin and VWF. Per control, 5 images were analyzed with the OrganelleProfiler pipeline. A) Graphical representation of the measurements that were performed on the objects. Eccentricity (top), length of Weibel-Palade bodies (WPBs) measured as maximum ferret diameter (middle) and distance of WPBs to the nuclei and the cell membrane was measured (bottom). Relative distance of the WPB to the nucleus in the cell was calculated as 100% x (distance to nucleus) / (distance to nucleus + distance to cell membrane). B) Cell count per image. C) The cell area (μm^2^) per cell of all 5 images pooled (n = 188 in group 1 and n = 52). D) Number of WPBs per cell. E) Distance of the WPB to the nucleus relative to their position in the cell in percentage. F) Mean WPB length per cell in μm. G) Mean eccentricity of WPBs per cell. Data is shown as median with min/max boxplot. Mann-Whitney test was performed on not normally distributed data (D and G). Unpaired T test with Welch’s correction was performed on normally distributed data (B, C, E and F); *p<0.05 **p<0.01, ***p<0.001.

#### Step VI – Quality control and analysis of output

For quality control, all objects’ outlines are overlaid on the VWF signal. This overlay allows the user to check whether the pipeline was accurate in the identification of objects. Cells are numbered so potential outliers can be easily identified and the pipeline can be adjusted if needed. The exported output can be used to quantify and perform qualitative analysis on images of interest.

Automated quantification using OrganelleProfiler revealed significant differences in cell count, cell area and number, size, shape and localization of WPBs between group 1 and group 3 ECFCs (Fig 3B-G). Data is shown as mean ± standard deviation (SD). Fig 3B shows a significantly lower number of cells per image in group 3 (10.40, ± 2.40) compared to group 1 (37.60, ± 2.80) (p=0.0003). Logically, as all ECFCs were confluent, we observed a larger mean cell area in group 3 (4016 μm^2^, ± 2445) than in group 1 (1143 μm^2^, ± 516.60) (Fig 3C) (p<0.0001). The total number of WPBs per image was lower in group 1 compared to group 3 (not shown). Additionally, the number of WPBs per cell was significantly lower in group 3 (30.92, ± 29.54) than in group 1 (107.30, ± 58.51) (p<0.0001) (Fig 3D). The distance of WPBs to the nuclei relative to their position in the cell was determined and shown in Fig 3E. The relative distance was significantly lower in group 3 ECFCs (32.31%, ± 23.62) when compared to group 1 (53%, ± 30.10) (p<0.0001) indicating that within the cell, WPBs were located closer to the nucleus in group 3 ECFCs. Finally, the mean WPB length was lower in group 3 (1.10 μm, ± 0.27) versus (1.38 μm, ± 0.21) in group 1 ECFCs (p<0.0001) and the WPBs were significantly more round in group 3 (0.63, ± 0.08)) versus (0.78 ± 0.04) (p<0.0001) (Fig 3F/G). The lower number of WPBs and the observation that they are smaller and rounder in group 3 when compared to group 1 could explain the decreased production and secretion of VWF observed previously (de Boer, JTH, 2020).

To further validate the quantitative data obtained from our automated OrganelleProfiler pipeline we also performed a manual quantification of several of these parameters using Fiji image analysis software, specifically the region of interest manager (12). One image of the group 1 ECFCs was used for the scoring. The manual scoring of the cells using the freehand selection resulted in 34 cells with a mean surface area of 1264 μm^2^, ± 497.93. For three cells all WPBs were scored using the straight line measuring the longest distance in the WPB. In these cells the manual scoring showed a mean WPB count 117, ± 38.63 and a length of 1.57 μm, ± 0,09. All measurements were compared with the CellProfiler measurements on the same image and none of the results differed significantly. Taken together, we can conclude that both measurements with CellProfiler and Fiji are comparable and thus CellProfiler can be used to accurately measure cells and organelles.

### OrganelleContentProfiler (OCP)- Automated measurement of proteins in secretory organelles

The OrganelleContentProfiler pipeline is an addition to the OrganelleProfiler pipeline. By adding 4 extra steps, secondary proteins of interest in, on or outside the organelle can be measured. For this purpose we analyzed the presence of Rab27A, a small GTPase that promotes WPB exocytosis and that is recruited to the WPB membrane during the maturation of these organelles after their separation from the Golgi complex (9, 10, 13). We also determined as a control the presence of protein disulfide isomerase (PDI), a marker for the endoplasmic reticulum which should not show specific localization in or on the WPBs (5, 14). Fig 4 shows example images of Rab27A as well as PDI co-staining in group 1 healthy donor ECFCs that were used in this pipeline. The CellProfiler modules that together form the OrganelleContentProfiler pipeline can be divided into 4 steps (Fig 5), which are described below. Further details on every module are described in S3 file.

**Fig 4.**
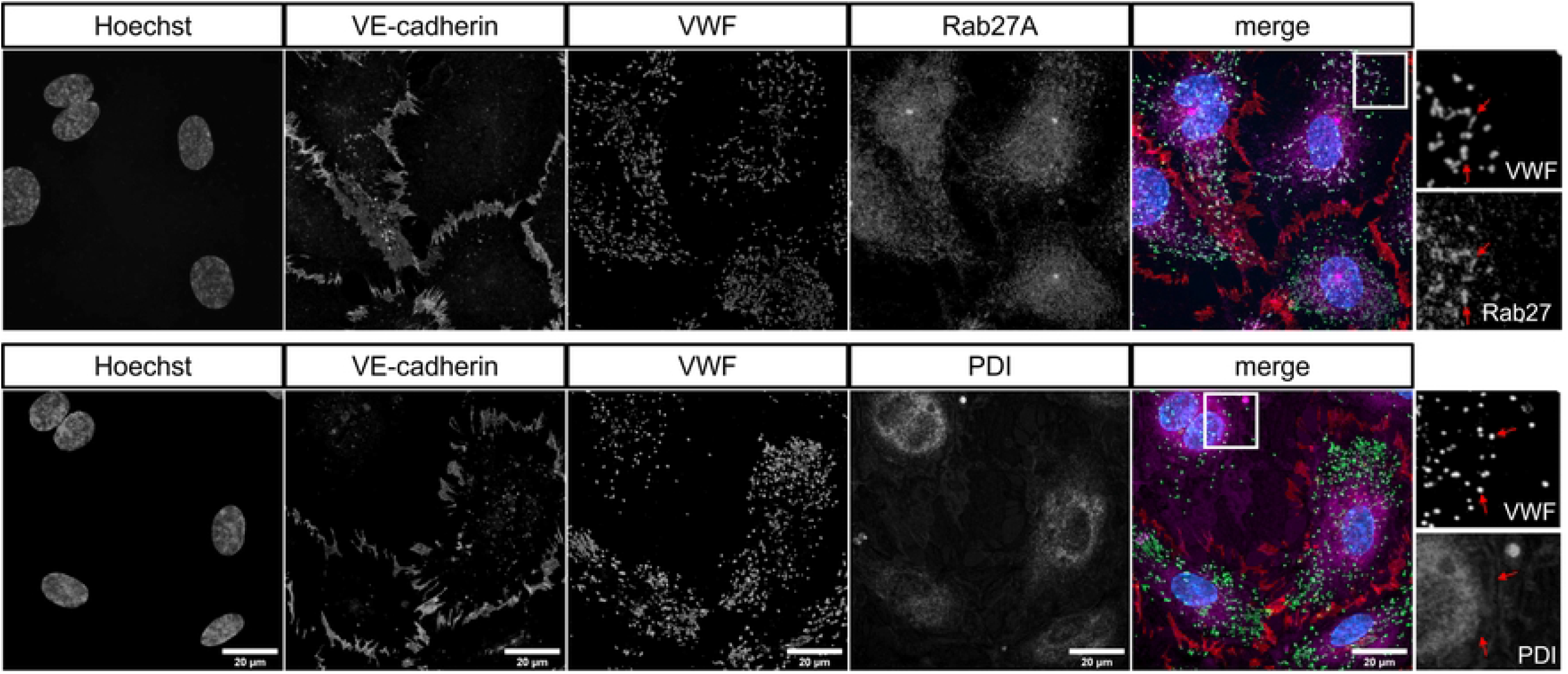
Representative images of one healthy group 1 ECFC control belonging to previously classified groups based on morphology (6). Cells were stained for Hoechst (blue), VE-cadherin (red), VWF (green) and Rab27A (top) or PDI (bottom). Scale bar represents 20 μm. Images were taken with a 63x objective. Red arrows indicate WPBs as identified in the VWF channel and the same location in the Rab27A or PDI channel.

**Fig 5.**
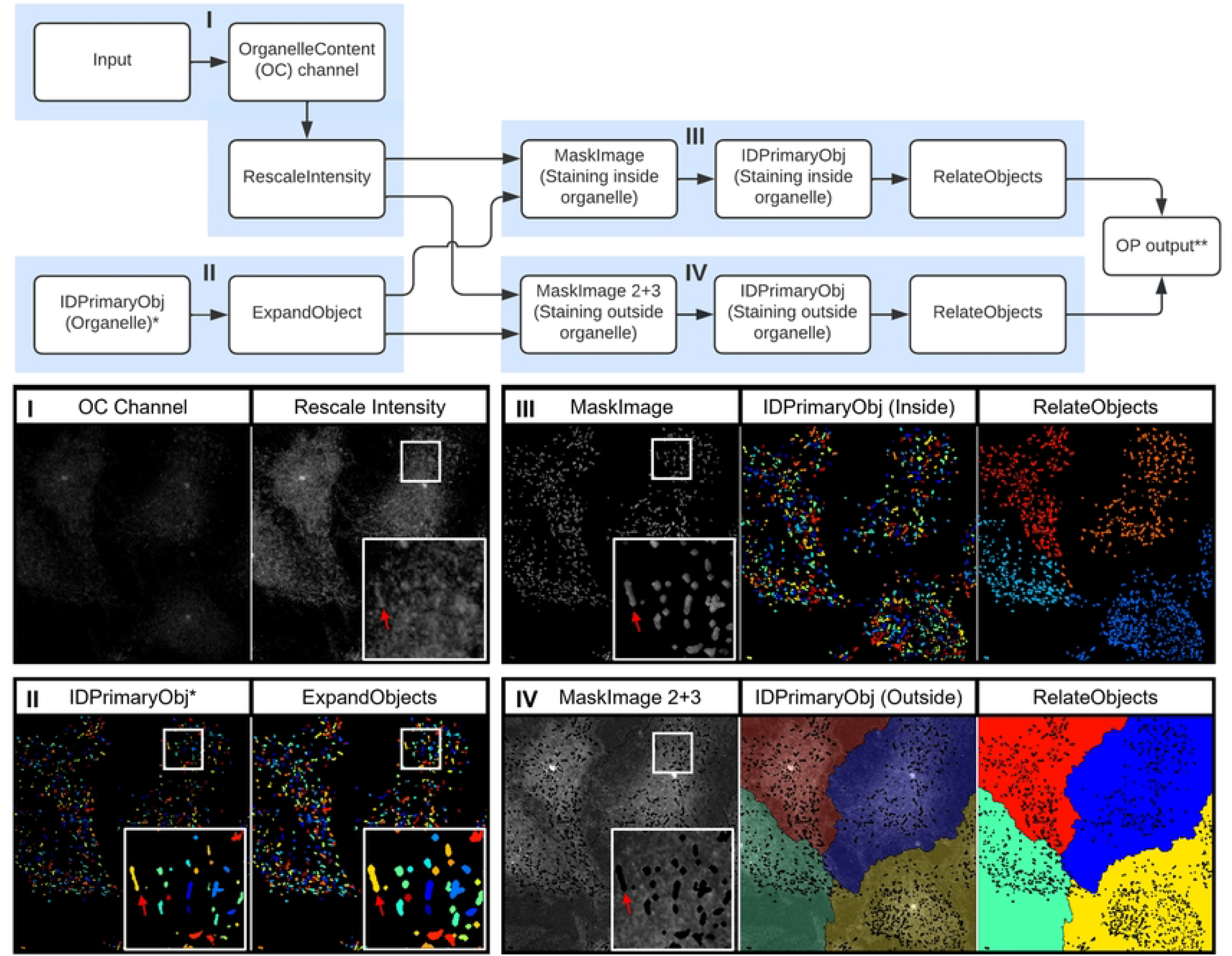
OrganelleContentProfiler: Quantitative and qualitative analysis of other organelle proteins. Top, flowchart of the modules within the OrganelleContentProfiler pipeline. I) Input of Rab27A (Organelle content) channel and rescaling of this channel. II) Input of primary object (Organelle) (left). and expansion of this object (right). III) Masking of the Rab27A channel using the Expanded organelle objects to leave only Rab27A signal inside the organelle (left). Identification of the Rab27A signal per WPB as object (middle) and relating these objects to the cells as child and parent respectively (right). IV) Masking of the Rab27A channel using the Expanded organelle objects to leave only Rab27A signal outside the organelle (left). Identification of the Rab27A signal in the cell as object without the WPBs(middle) and relating these objects to the cells as child and parent respectively (right). * Identified in step IV of the OrganelleProfiler pipeline (Fig 2). ** Pipeline continues with step V and VI from the OrganelleProfiler pipeline.

#### Step I – Input of an additional channel

Similarly to the OrganelleProfiler, images are imported into the software. In this example, images have one additional channel containing the staining for either Rab27A or PDI. Again, channels are separated and the fourth channel is rescaled in order to view the channel in the final quality control.

#### Step II – Import of organelle object identified in the OrganelleProfiler pipeline

In this step, the organelle object as identified in the OrganelleProfiler pipeline is modified. The objects are initially identified using the staining for VWF, which is a cargo protein that is contained within the organelle. The secondary protein of interest, Rab27A, is a membrane protein that is located on the cytoplasmic face of the WPB membrane. To ensure full encapsulation of the Rab27A signal the object is therefore expanded by 2 pixels in all directions.

#### Step III and IV – Identification of the secondary protein of interest “inside” and outside the organelle

In these parallel steps, the expanded organelle objects and the rescaled secondary staining channel are used. The expanded objects are used as a mask to remove all signal of the Rab27A or PDI staining outside the organelle (step III) and inside the organelle (step IV). The remaining signal is then identified as object, resulting in two new objects containing the signal inside the organelles and outside the organelles respectively. These new objects are processed according to step V from the OrganelleProfiler including the measurements, relating, quality control and export.

#### Step V and VI – Measurements, quality control and analysis of results

In the OrganelleContentProfiler pipeline, different stainings on the same ECFC control are compared. In addition to the output from the OrganelleProfiler, the OrganelleContentProfiler provides measurements of the intensity of a secondary signal inside the organelle. Furthermore, it can quantify the cytoplasmic intensity values outside of the organelle which can be used for correction of the “inside” organelle signal. Signal intensity is noted as arbitrary intensity units (A.U.) as microscopes are not calibrated to an absolute scale.

We first confirmed that the number of WPBs quantified using OrganelleContentProfiler does not depend on the co-staining used (Rab27A: 204.8 ± 70.48; PDI: 146, ± 56.38; p=0.24) (Fig 6A). ECFCs were stained with Hoechst and with antibodies against VE-cadherin, VWF and Rab27A or PDI. Fig 6B shows the A.U. inside and outside organelles and the A.U. inside the organelle corrected for the outside value. First, the VWF A.U. was analyzed as a measurement of a protein that is located predominantly in the WPB. The results show that the VWF A.U. values outside the WPBs was nearly zero (0.00076 ± 0.00033) and differed significantly from the inside A.U. (0.028 ± 0.0040) (p=0.0016) indicating that VWF is almost exclusively present in WPBs. Secondly, it was determined that the Rab27A staining shows a significantly higher A.U. inside (0.081 ± 0.0085) the WPBs when compared to the outside measurement (0.052 ± 0.0041) (p=0.0062). From this it can be concluded that part of the Rab27A protein is present in or on the WPB. Finally, the A.U. of the PDI staining was analyzed. PDI is only present inside the endoplasmic reticulum and should not yield increased A.U. inside the WPB. Indeed, the A.U. inside (0.074 ± 0.016) and outside (0.062 ± 0.0082) the organelle were similar (p=0.20), indicating that PDI is not located specifically in or on WPBs.

**Fig 6.**
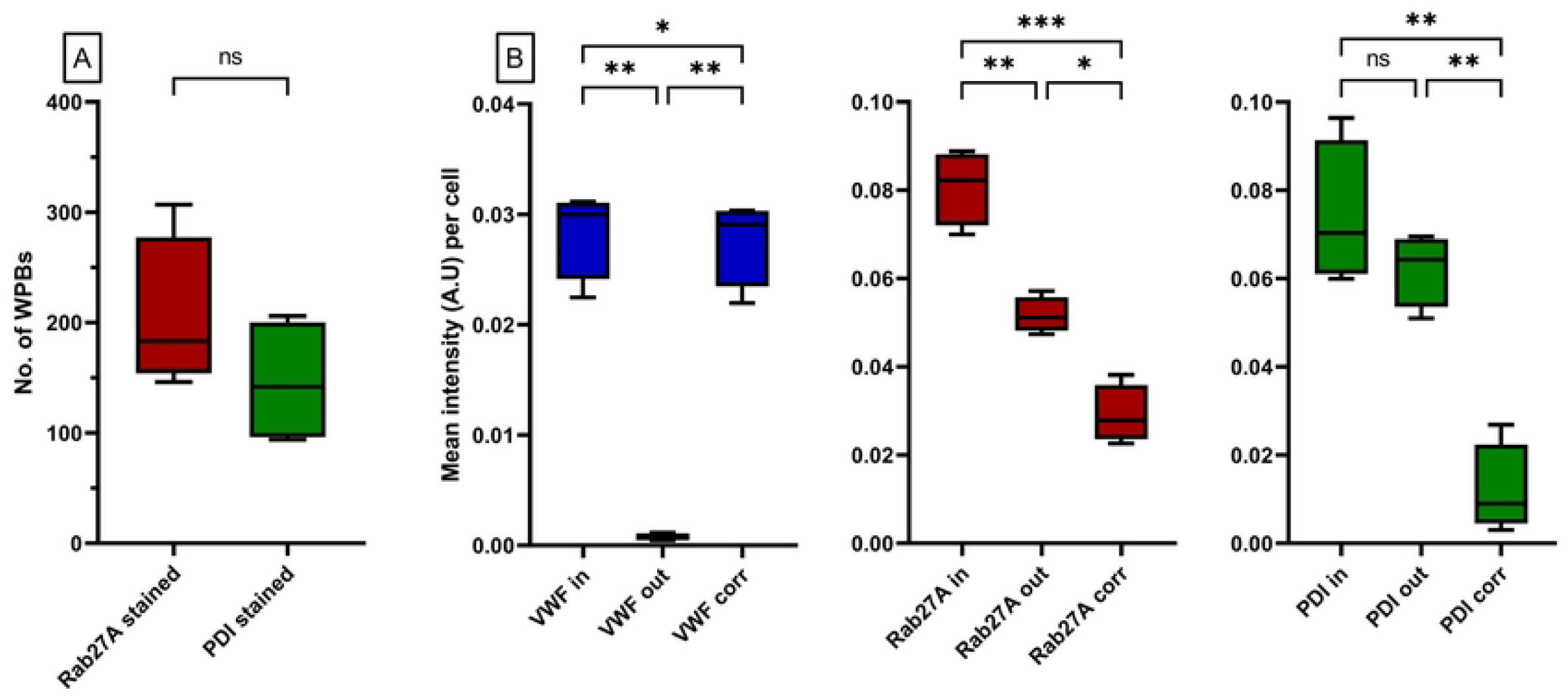
Quantification of signal intensity inside cell organells. A group 1 ECFC control as defined previously (6) was stained with Hoechst and with antibodies against VE-cadherin, VWF and PDI or Rab27A. One image per staining was analyzed using the OrganelleContentProfiler pipeline. A) Both images had the same number of cells (n=4) and the same number of WPBs. B) the mean intensity in arbitrary intensity units (A.U.) per cell. for the PDI (left), Rab27A (middle) and VWF (right) staining. Each graph shows the measured mean intensity inside (in) the WPBs, outside (out) the WPBs and the intensity inside the WPB after correcting for the out signal (corr). Data is shown as median with min/max boxplot. RM one way ANOVA was performed with Geisser-Greenhouse correction; *p<0.05 **p<0.01, ***p<0.001.

Once more, to validate the quantitative data obtained by CellProfiler, we also performed a manual scoring using Fiji of the A.U. for the Rab27A and VWF staining inside and outside of all WPBs (n=199) in one cell. We observed that the results determined manually using Fiji (A.U. VWF inside = 0.039, VWF outside = 0.000029; Rab27A inside = 0.094, Rab27A outside = 0.053) lie within the same range as those determined by CellProfiler. This shows that the OrganelleContentProfiler can determine organelle specific stainings and measure the intensity of the staining corrected for the cytoplasmic value.

## Discussion

Quantifying large numbers of organelles is challenging due to the density and morphological heterogeneity of the organelles. The pipelines described here can be used to overcome these challenges and can provide organelle analysis in great detail on a larger scale. The OrganelleProfiler allows for measurement of cell and nucleus quantity and shape, and organelle quantity, shape, size and location within the cell. The organelles are also related to the cells which allow for cell-by-cell analysis. This information can be used to determine differences between a heterogeneous cell population or between patient and control cells. The OrganelleProfiler pipeline has shown significant differences between group 1 and group 3 ECFC controls based on only 5 images. Once optimized for a set of images, the pipeline can analyze thousands of cells and hundreds of thousands of organelles within hours without potential bias associated with manual image processing and quantification. Furthermore, with the OrganelleContentProfiler, secondary organelle markers can be measured and quantified. We showed 3 stainings of proteins with different localizations; PDI, which is only present on the endoplasmic reticulum, Rab27A which is present in the cytoplasm, but is also trafficked to the WPBs, and VWF which is mostly present in WPBs. Using the OrganelleContentProfiler pipeline we were able to quantify these stainings and determine the localization of these proteins. It is also possible to measure other organelle stainings at the same time by duplicating modules 3 to 8 of this pipeline and adjusting these for the additional channels.

Finetuning of the smoothing, thresholds and enhancement of the signal is necessary to ensure correct identification of objects. For every image set, a balance must be found to prevent over and under segmentation of organelles. Despite optimization, perfect segmentation of organelles is not always possible, especially in areas where organelles are crowded together. These imperfections may lead to incorrect identification of organelles, which could play out as underestimations of WPB numbers or overestimation of WPB dimensions. However, as all images are analyzed by the same pipeline, this error is expected to occur to a similar extent in all samples. One point of improvement on the OrganelleContentProfiler pipeline could be the correction of the organelle secondary staining with the intensity levels directly surrounding the organelle instead of the mean intensity in the entire cytoplasm. This was not possible within the CellProfiler software but could be done in data processing afterwards using the MeasureObjectIntensityDistribution module and relating this to the distance of WPBs to the nucleus (7).

A comparison with manual scoring using Fiji was performed to check the validity of the results generated by our automated pipelines. We generally found that the results obtained with OrganelleProfiler and OrganelleContentProfiler correspond very well with manual quantifications using Fiji, although subtle differences were found for two parameters. First, the maximum ferret diameter is calculated based on the smallest convex hull that is created around the WPB. The manual scoring measured the length of the WPB in a line and not as the inside of a convex hull. This could cause the slight difference in length as measured between CellProfiler and Fiji. Second, VWF and Rab27A intensities inside WPBs as determined by manual scoring was slightly higher than from the OrganelleContentProfiler measurements. Possibly, the outlines that were drawn around WPBs manually were more strict than those generated by OrganelleContentProfiler, because the human eye is less capable at detecting the very small changes in signal intensity near the edges of the organelles. As such, the signal intensities in these edges may have not been included in the manual analysis, resulting in a higher mean value per WPB.

To conclude, the OrganelleProfiler and OrganelleContentProfiler pipelines provide powerful, high-processing quantitative tools for analysis of cell and organelle count, size, shape, location and content. These pipelines were created with the purpose of analyzing morphometric parameters of WPBs in endothelial cells, but they can be easily adjusted for use on different cell types or organelles. This can be especially useful for analysis of large datasets where manual quantification of organelle parameters would be unfeasible.

## Abbreviations

OP: OrganelleProfiler
OCP: OrganelleContentProfiler
ECFC: Endothelial colony forming cells
WPB: Weibel-Palade body
VWF: Von Willebrand factor
VWD: Von Willebrand disease
A.U.: Arbitrary intensity units

## Acknowledgements

The SYMPHONY consortium, which aims to orchestrate personalized treatment in patients with bleeding disorders, is a unique collaboration between patients, health care professionals, and translational and fundamental researchers specializing in inherited bleeding disorders, as well as experts from multiple disciplines. It aims to identify best treatment choice for each individual based on bleeding phenotype. To achieve this goal, work packages (WP) have been organized according to 3 themes (e.g. Diagnostics [WPs 3 and 4], Treatment [WPs 5-9], and Fundamental Research [WPs 10-12]). Principal investigator: M.H. Cnossen; project manager: S.H. Reitsma.

## Authorship Contributions

SNJL and RJD performed research, developed quantification methods and analyzed data; SNJL, JE, and RB designed the research and wrote the paper.

## Supporting information

**S1 table. Supporting information on antibodies**

**S1 file. OrganelleProfiler pipeline**

**S2 file. OrganelleContentProfiler pipeline**

**S3 file. Detailed guide per module of the OP and OCP pipelines**

